# “Centrosome Amplification promotes cell invasion via cell-cell contact disruption and Rap-1 activation”

**DOI:** 10.1101/2022.05.09.490051

**Authors:** Anu Prakash, Shishir Paunikar, Mark Webber, Emma McDermott, Sri H. Vellanki, Kerry Thompson, Peter Dockery, Hanne Jahns, James A.L. Brown, Ann M. Hopkins, Emer Bourke

## Abstract

Centrosome amplification (CA) is a prominent feature of human cancers linked to genomic instability and tumourigenesis *in vivo*. CA is observed as early as pre-malignant metaplasia, with CA incidence increasing as the disease progresses from dysplasia to neoplasia. However, the mechanistic contributions of CA to tumourigenesis (tumour architecture and remodelling) are poorly understood.

Using non-tumourigenic breast cells (MCF10A), we demonstrate that CA induction (by CDK1 inhibition or PLK4 overexpression) alone increased both cell migration, invasion and Extracellular Matrix (ECM) remodeling. Mechanistically, CA induction activated small GTPase Rap-1. We demonstrated the key role of Rap-1 mediated signalling in CA induced tumourigenesis through Rap-1 inhibition (using GGTI-298) which blocked CA-induced migration, invasion and ECM attachment.

CA induction in a long-term MCF10A cell culture system disrupted epithelial cell-cell junction integrity, via dysregulation of expression and subcellular localisation of cell junction proteins (ZO-1, Occludin, JAM-A & β-catenin). At the ultrastructural level, CA significantly inhibited apical junctional complex formation, as visualized by transmission electron microscopy. CA induction in the luminal A breast cancer cell line MCF7 revealed similar trends in cell junction disruption. Furthermore, CA induction in MCF10A elevated expression of integrin β-3, matrix metalloprotease MMP1 and MMP13 facilitating the observed ECM attachment, degradation and cell invasion phenotype.

*In vivo* validation using a Chicken Embryo xenograft model, showed CA positive (CA+) MCF10A cells invaded into the chicken mesodermal layer, characterised by inflammatory cell infiltration and a marked focal reaction between chorioallantoic membrane and cell graft. This reaction was inhibited by pre-treatment of CA+ MCF10A cells with GGTI-298. Interestingly, in metastatic breast cancer cells with high levels of endogenous CA (triple negative cell line MDA-MB-231) inhibition of this CA-signalling pathway (using PLK4 inhibitor Centrinone B) abrogated their metastatic capacity *in vitro*. This demonstrates dual roles for CA signalling, for initiating and maintaining the CA-induced metastatic phenotype.

Here, we demonstrated that CA induction in normal non-tumourigenic cells acts through Rap-1-dependent signaling to confer early pro-tumourigenic changes promoting tumour progression, mediated by ECM disruption, and altered cell-cell contacts. These insights reveal that in normal cells, CA induction alone (without additional pro-tumorigenic alterations) is sufficient to induce tumourigenesis and CA-mediated signaling supports a metastatic phenotype.

**Statement:** Centrosome amplification alone drives early tumourigenic change in normal breast epithelial cells

## INTRODUCTION

Centrosomes play essential roles in averting genomic instability by regulating cell cycle control, spindle assembly and ultimately cell replication. Centrosome amplification (CA), a numerical centrosomal defect (≥3 centrosomes per cell) is a hallmark feature of many high-grade human tumours (Chan, 2011). CA is triggered by well-characterised mechanisms including cytokinesis failure, dysregulation of centrosome duplication proteins and pericentriolar material (PCM) fragmentation (reviewed in (Godinho *et al*, 2014)). Additionally, extra centrosomes are directly induced *in vitro* by DNA damage and defects in DNA repair pathways linking CA to inherent genetic instability of tumour cells (Bourke *et al*, 2010; Bourke *et al*, 2007; Dodson *et al*, 2004). However, it is increasingly evident that aberrant centrosome numbers are not necessarily a downstream consequence of tumorigenesis and centrosome abnormalities directly perpetuate genomic instability by promoting chromosome missegregation (Cosenza & Kramer, 2016; Ganem *et al*, 2009) and cellular invasion via cell-autonomous and non-autonomous mechanisms (Adams *et al*, 2021; Arnandis *et al*, 2018). CA induction promotes tumourigenesis *in vivo* (Coelho *et al*, 2015; Levine *et al*, 2017; Sercin *et al*, 2016) and has been observed in pre-malignant metaplasia, with CA rates increasing concurrently with disease progression to dysplasia, and is often accompanied by p53 mutation and loss (Lopes *et al*, 2018; Pihan *et al*, 2003; Segat *et al*, 2010). Together, this evidence suggests that CA plays a crucial role in early tumour initiation and malignant progression.

In its capacity as the major microtubule organising centre (MTOC), the centrosome orchestrates microtubule (MT) assembly pending mitosis while also controlling cell positioning, polarity and motility through its links to the acto-myosin cytoskeleton, cell-cell and cell-extracellular matrix (ECM) junctions (Bettencourt-Dias, 2013; Bornens, 2012). Cell-cell attachments are key to maintaining barrier function in polarised epithelia, and are mediated by highly specialised junctional complexes consisting of tight junctions, adherens junctions, desmosomes and gap junctions (Garcia *et al*, 2018). Crucially, the formation and regulation of these junctions is dependent on their interactions with cytoskeletal polymers of microtubules and actin, pools of which can nucleate from the centrosome (Farina *et al*, 2016; Farina *et al*, 2019; Vasileva & Citi, 2018). Therefore abnormalities of the centrosome, structural or numerical, may potentially impact on cell-cell junction function, promoting cell detachment from the epithelial layer. Indeed, recent studies have shown that structural centrosome anomalies induced by ninein-like protein (NLP) overexpression in MCF10A breast epithelia, disrupt epithelial polarity and cell-cell contacts via dysregulation of E-cadherin localisation within adherens junctions mediated by small GTPase Rac1 and Arp2/3 signalling (Ganier *et al*, 2018). This leads to increased cell stiffness, detachment from the epithelium and cell budding towards the ECM. CA induced by PLK4 overexpression in MCF10A cells, generates increased MT nucleation and polymerisation, also causing Rac1 activation and Arp2/3-dependent actin polymerisation, leading to invasive lamelipodia formation in 3D cultures. The invasive phenotype is accompanied by loss of cell-cell adhesion, disruption of junction size and positioning as well as dysregulation of E-cadherin junctional localization (Godinho *et al*., 2014). In human umbilical vein endothelial cells (HUVEC), CA induction alters cell polarity and perturbs luminogenesis of newly formed endothelial sprouts accompanied by chaotic localization of vascular endothelial (VE)-cadherin, resulting in the disruption of adherens junction stability (Buglak *et al*, 2020). While data suggests the CA-induced metastatic phenotype is mediated by perturbations of cell-cell and cell-ECM contacts, clear evidence of the mechanism remains to be identified and validated.

Another central step towards local invasion of tumour cells into adjacent tissue is the degradation of ECM components via matrix metalloproteinase (MMP) secretion (Kessenbrock *et al*, 2010). This family of zinc-dependent endopeptidases are involved in physiological tissue remodelling as well as pathological ECM breakdown within the tumour microenvironment. Emerging evidence of the role of CA in cell migration and invasion points to a possible link between CA and increased MMP expression. *In vivo*, tumours derived from the injection of p53-null tetraploid cells into mice are associated with high levels of centrosome amplification, accompanied by amplification of an MMP gene cluster (Fujiwara *et al*, 2005). Furthermore, treatment with the broad spectrum matrix metalloprotease (MMP) inhibitor, marimastat (BB-2516), decreased the fraction of CA-induced invasive acini in MCF10A 3D cultures (Godinho *et al*., 2014). Proposed intracellular mediators of CA-induced effects include the subfamilies of small GTPases, Rho and Ras, which play roles as molecular switches integrating signals from the membrane to actin-cytoskeletal changes to a wide range of effector proteins of migration and invasion, including the expression of MMPs (Sahai *et al*, 2001). It has been shown that the Rho-GTPase Rac1 is activated by CA induction in breast epithelial cells and that a Rac1 inhibitor (NSC23766) partially rescued the invasive CA-induced phenotype (Godinho *et al*., 2014). However, the complex and multifaceted range of signalling pathways activated by CA-induction, which facilitates cellular invasion and ECM remodelling, requires further detailed characterisation.

In this study, we investigate how CA induction alone (in “normal” non-tumourigenic cells) can drive the initial stages of tumourigenesis by promoting cell invasiveness (in both non-tumourigenic and tumourigenic breast epithelial cells) via alteration of extracellular contacts (cell-cell and cell-ECM) and dysregulated MMP/TIMP production, mediated through Rap1 signalling. The results were validated using both an *in vitro* semi-3D long-term cell culture system, and an *in vivo* (chick embryo xenograft) model. *In vivo* implantation of CA-induced (+CA) MCF10A cells resulted in cell invasion into the chorioallantoic membrane and a vigorous proliferative focal reaction involving immune cell infiltration.

## RESULTS

### CA increases cellular migration and invasion *in vitro* and in an *in ovo* chick embryo xenograft model

Polo-like kinase (PLK4) is the master regulator kinase of the centrosome duplication cycle (Bettencourt-Dias *et al*, 2005; Habedanck *et al*, 2005). Non-tumourigenic breast epithelial MCF10A PLK4 cell lines employed in the current study were engineered with doxycycline (Dox)-inducible PLK4 transgenes. Previous studies using these cell lines show that Dox-induced Wild Type PLK4 expression induces CA (CA+ MCF10A) while a Dox-induced truncated PLK4 variant PLK4^1-608^ (lacking the C-terminal centrosome localization domain; amino acids 1–608) does not induce CA (Godinho *et al*., 2014). Here, Dox treatment of MCF10A PLK4 cells (CA+ MCF10A) led to a significant (p≤ 0.02) CA induction in 80% (±1.26 SEM) of cells after 48h (Fig. 1A-B). We measured cell invasion and migration post-CA induction using Matrigel-coated and uncoated transwell assays respectively. CA+ MCF10A PLK4 cells showed a significant (p≤ 0.05) ~3-fold increase in invasion (Fig. 1C), and a significant (p≤ 0.02) ~2-fold increase in migration (Fig. 1D), compared to both untreated controls and PLK4^**1-608**^. To independently validate that these findings are dependent on CA-induction, we induced CA in wild-type MCF10A by CDK1 inhibition, as previously demonstrated (Bourke *et al*., 2010), using the CDK1 pharmacological inhibitor RO3306 (Steere *et al*, 2011). Confirming previous reports, RO3306 treatment significantly induced CA (p=0.2), migration and invasion (p≤ 0.05) (Fig. S1A-C).

**Figure 1:**
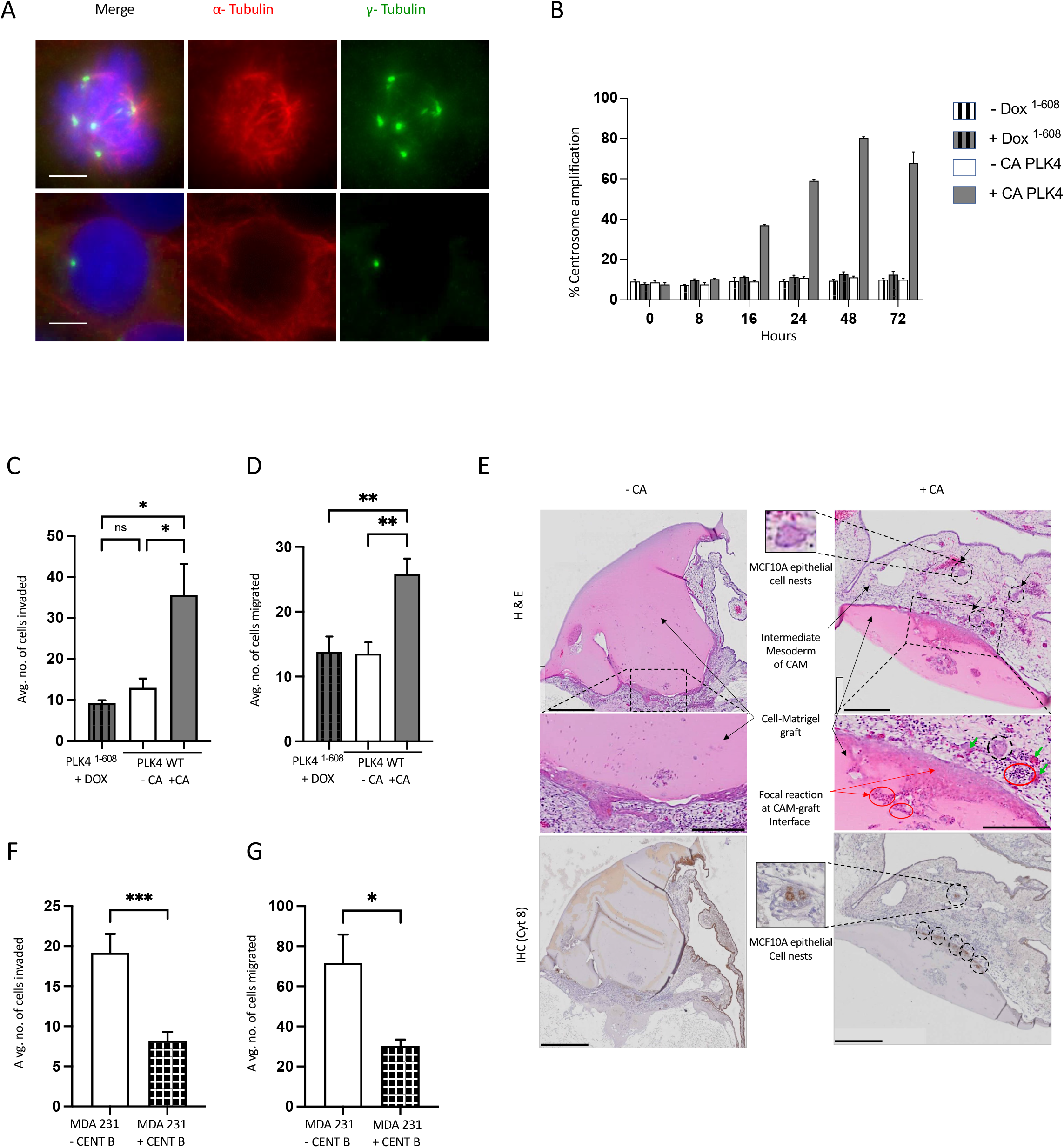
CA increases *in vitro* and *in vivo* migration and invasion and blocking CA decreases metastatic characteristics of MDA-MB-231. (A) Representative images of immunofluorescently stained MCF10A cells fixed with Methanol/5mM EGTA solution and stained for microtubules (α-tubulin, red), centrosomes (γ-Tubulin, green) and counterstained with DAPI (DNA, blue). Scale bar = 10 μm. Images are representative of n=3 independent experiments. (B) Fluorescence microscopy quantification of % of CA+ cells at each time point post-Dox treatment; normal centrosome number (≤2 per cell) and amplified centrosomes or CA (≥3 per cell). Bar graphs represent mean ± SEM from 3 independent experiments; ≥200 cells/time point. (C) Transwell invasion and (D) migration assay results show that CA induction significantly increased cellular invasion and migration in MCF10A PLK4 cells compared to controls CA-PLK4 and DOX+ PLK4^**1-608**^. The bar graph represents mean (± SEM) number of cells invaded and migrated from four independent experiments. One-way ANOVA and Bonferroni’s multiple comparison test was used to determine significance, *P ≤ 0.05, **P ≤ 0.01. (E) Representative images of chicken xenograft assay with MCF10A +/- CA cell-matrigel graft. Images show Haemotoxylin & Eosin (H&E upper panels) (matrigel staining pink) and immunohistochemical staining for Cytokeratin 8 (Cy8: lower panels) on formalin-fixed paraffin-embedded tissue sections CA± MCF10A PLK4 cells/matrigel grafts. The CA+ graft shows moderate to strong chicken-graft focal reaction (red arrows), with multifocal heterophil infiltration in the intermediate mesoderm layer (red circles) and invasion of epithelial cell nests into chicken mesodermal layer (black arrows/circles). Mild to moderate focal reaction of the CAM to the graft was noted in CA-grafts. The corresponding IHC (lower panels) showed that the invaded epithelial cell nests were positive for Cy8 (dotted circles). Scale bars = 500μm/250 μm. (F) Transwell invasion and (G) migration assays show that blocking endogenous CA with 150nM Centrinone B for 16 hours in TNBC cell line MDA-MB-23 significantly decreased cellular invasion and migration. Migrated or invaded cells were quantified with DAPI staining and the cells counted using IF from five different fields at 20X magnification. Bars represent mean ± SEM values from two independent biological repeats and an unpaired two-tailed Student T-test was used to determine the statistical significance: *P ≤ 0.05, ***P < 0.001.

To investigate how CA induction could initiate early tumourigenic change *in vivo*, we employed a chicken embryo xenograft model. The Chorioallantoic membrane assay (CAM assay), is a rapid, cost effective *in vivo* model for the study of tumourigenesis, angiogenesis, invasion and metastases (reviewed in (DeBord, 2018; Komatsu *et al*, 2019; Miebach *et al*, 2022; Ribatti, 2014). The CAM assay has been used to study angiogenesis and tumour cell invasion in breast, pancreatic, colorectal, prostate, brain and ovarian cancers using both human tumour cell line-derived xenografts as well as patient-derived xenografts (Aaltonen *et al*, 2022; Harper *et al*, 2021; Lokman *et al*, 2012). The highly vascularised chicken chorioallantoic membrane of the fertilised chicken egg (onto which the MCF10A cells are grafted) consists of the chorionic epithelium (the outer layer) with a superficial vascular plexus, and the intermediate mesoderm layer and the allantoic epithelium (inner layer). The CAM assay allows characterisation of extremely early changes post-CA induction, not previously observable using *in vivo* mouse CA models (Coelho *et al*., 2015; Levine *et al*., 2017; Sercin *et al*., 2016)). Matrigel embedded Control or CA+ MCF10A cells were implanted onto the chick chorioallantoic membrane and allowed to grow for 5 days. Histological evaluation showed that the CA+ MCF10A PLK4 invaded into the chorionic epithelium (CE) (at the cell-matrigel interface) (black arrows, Fig. 1E) inducing a multifocal reaction (ranging from moderate to marked) of the CE including proliferation of mesodermal cells which spread extensively into graft area (red arrows, Fig. 1E), multifocal heterophil infiltration (red circled area, Fig. 1E), and dilated lymphatics in the intermediate mesoderm layer (green arrows, Fig. 1E). Additional representative images of matrigel:CE reactions are shown in Fig. S2 A-C. CA+ MCF10A cells (labelled with anti-human Cytokeratin 8) invaded into the chicken tissue, forming epithelial cell nests within the intermediate chicken mesodermal layer (Fig. 1E dotted circles and enlarged panel). Quantification of CAM invasion into the matrigel-cell graft showed that CA increased the area of invasion significantly compared to CA- grafts (p≤ 0.001, Fig. 2F). Interestingly, histological analysis indicated that CA+ MCF10A sections displayed clear secondary tumour characteristics including haemorrhaging, necrosis and calcification at the CAM/matrigel interface (Fig. S3 A-C). Together these results show that CA induction alone in non-tumourigenic cells is sufficient to induce a migratory and invasive cellular phenotype (both *in vitro* and *in vivo*), and induction of secondary tumour-associated changes in the surrounding tissue *in vivo*.

**Figure 2:**
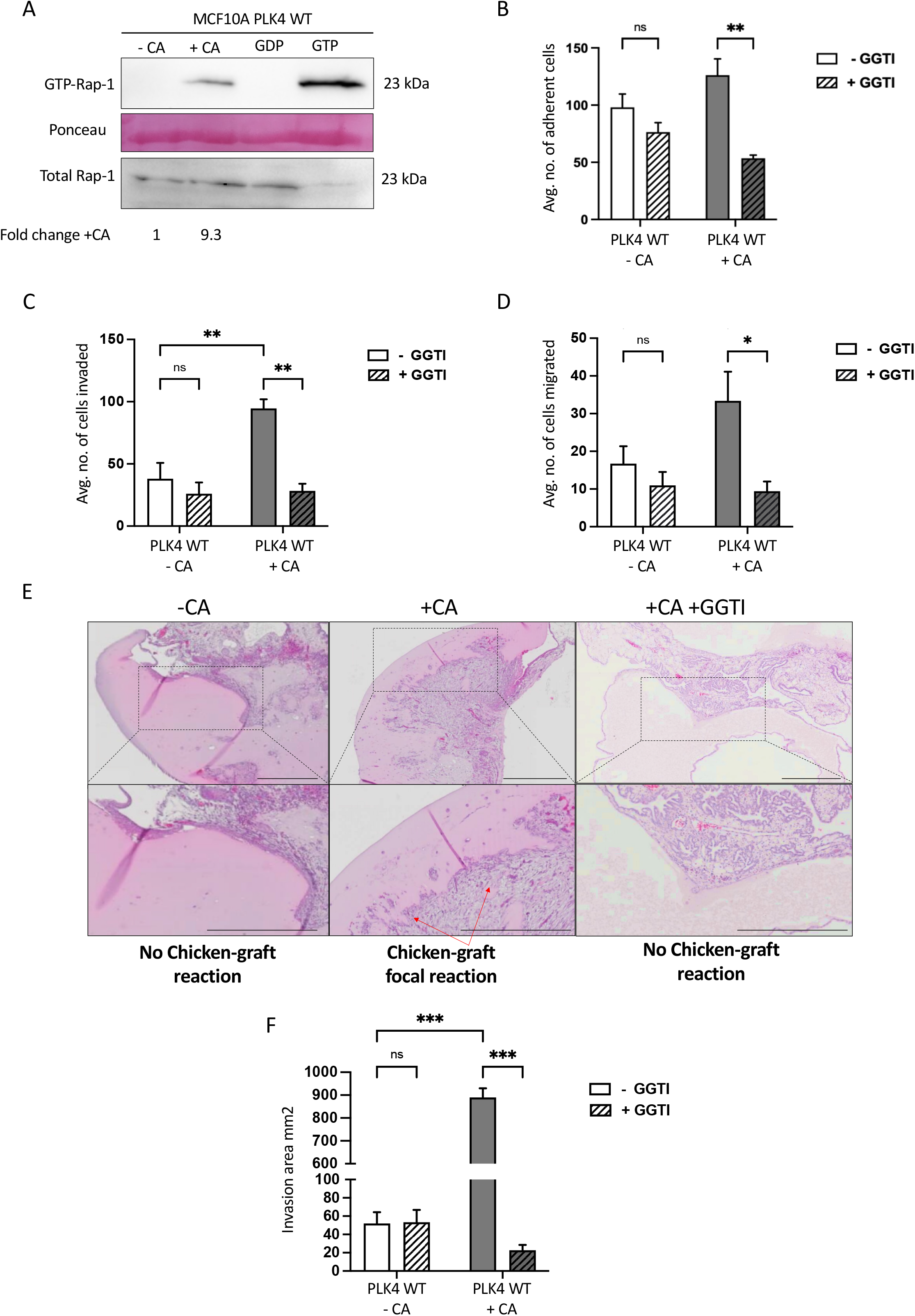
CA induced migration, invasion and cell-ECM attachment involves activation of Rap1. (A) Western blot of Rap1 pull-down assay in MCF10A PLK4 shows CA-induced increased GTP-bound Rap-1. Negative (GDP) and positive (GTPγS) controls on untreated cell lysates were included. Ponseau stain is included as a loading control for the pull-down. Total Rap-1 in 60μg of total cell lysates is shown in the bottom panel. The image shown is representative of n=3 independent repeats and the fold-change represents the quantification of CA± bands by densitometry (B) CA increases MCF10A cell-ECM adherence in a Rap1-dependent manner. Cell adhesion assays was performed on MCF10A PLK4 cells 48h post-dox +/- 2 μg/ml treatment and 3h treatment +/- 10μM GGTI-298. Cells bound to the matrix were quantified with crystal violet staining-five 20X fields per well were counted and the bar graph represents mean ± SEM from four independent experiments. Two-way ANOVA was used to determine statistical significance, **P ≤ 0.01. (C-D) Rap1 inhibition significantly decreased CA-induced cellular invasion and migration. The bar graphs represent mean number of cells invaded and migrated, quantified by crystal violet staining in five random 20X fields per insert, ± SEM from four independent experiments. Two-way ANOVA and Bonferroni’s multiple comparison test was used to determine the significance, *P ≤ 0.05, **P ≤ 0.01. (E) Representative H&E images of chicken xenograft assay with cell-matrigel graft. MCF10A PLK4 cells were xenografted 48h post-dox (+/- 2 μg/ml) treatment and 3h treatment +/- 10μM GGTI-298. Rap-1 inhibition decreased the +CA-induced bidirectional chicken-graft focal reaction *in vivo*. Scale bar = 500μm. (F) Area (mm2) of chicken-graft focal reaction was quantified in scans of MCF10A grafts ± CA ± GGTI-298 using Olympus OlyVIA software and bar graphs represent mean ± SEM from three independent experiments. Two-way ANOVA and Bonferroni’s multiple comparison test were used to determine the significance, ***P ≤ 0.001.

To evaluate how CA contributes to an invasive phenotype, we utilized metastatic MDA-MB-231 cells (triple negative breast cancer; TNBC) which have endogenously high CA levels. CA was blocked using the PLK4 specific inhibitor Centrinone B. <48 hours Centrinone B treatment has been previously reported to reduce centrosome and centriole numbers, reverting CA lines to a normal centrosome complement (1-2 centrosomes), with >48 hours treatment leading to complete centrosome depletion (Denu *et al*, 2020; Denu *et al*, 2018; Marteil *et al*, 2018; Wong *et al*, 2015). Centrinone B treated MDA-MB-231 had CA reduced significantly (p≤ 0.05) from 39.75 (± 4.6%) to 10.25 (± 0.35%) (data not shown). Furthermore, Centrinone B treated MDA-MB-231 cells showed significantly (p≤ 0.001) reduced invasion (2.5-fold), and significantly (p≤ 0.05) reduced migration (2.7-fold) (Fig. 1F-G).

To further explore the observed Centrinone B effects on migration or invasion a second invasive TNBC cell line, MDA-MB-468 cells, with low endogenous CA levels (5-10% cells, data not shown) was tested. Interestingly, Centrinone B treatment of MDA-MB-468 cells did not significantly reduce migration or invasion (a reduction of 3% and 16% respectively; Fig. S1D). This suggests that in cancer cells with high CA, the increased migration and invasion is dependent on increased CA signaling. However, in cancer cells with low CA, the increased migration and invasion is independent of CA.

### CA induced migration and invasion involves activation of Rap1

It has been reported that activation of Rap1 (from inactive GDP-bound to active GTP-bound) can cause increased metastasis in cancer cells, playing a key role in regulating cell-ECM contacts (Sit & Manser, 2011). An activated Rap1 pull-down assay revealed that CA+ MCF10A increased Rap1 activation (Fig. 2A).

Supporting this, CA increased cell-ECM attachment (in comparison to untreated cells, Fig. 2B). Demonstrating the importance of Rap1 activation in CA-induced ECM remodelling, Rap1 inhibitor (GGTI-298) significantly (p ≤ 0.01) decreased this CA-induced cell-ECM attachment (Fig. 2B). Also, GGTI-298 treatment significantly decreased both the CA-induced cell invasion and migration (Fig. 2C&D; transwell assays). Furthermore, in the chick xenograft model, MCF10A PLK4 +CA cells treated with Rap1 inhibitor prior to engraftment showed a significant (p≤ 0.001) decrease in the CA-induced bidirectional chicken-graft focal reaction *in vivo* (Fig. 2E-F).

### CA induced disruption of epithelial barrier integrity

Cell adhesiveness is dysregulated in tumours, resulting in defective and altered histological structure promoting cancer invasion and metastasis (Hirohashi & Kanai, 2003). The effects of CA on epithelial barrier integrity and apical junction complexes are unknown. We explored the effects of CA on cell-cell adhesion and cell-ECM adhesion. Under standard cell culture conditions, MCF10A form poor tight junctions (TJ) (Fogg *et al*, 2005; Underwood *et al*, 2006). Therefore, to study the effect of CA on apical junctional complexes (AJC), an established long-term MCF10A cell culture assay which forms robust TJ, was used (Fig. 3A) (Marshall *et al*, 2009). Longterm CA-induced cultures, showed a significant increase (p ≤ 0.001) in CA+ MCF10A cells at the apical (42%) and middle (25%) regions of the multi-layered, apico-basally polarised cell culture system (Fig 3B-C). CA induced disruption of β-catenin staining from well-ordered cortical rings to a disordered incomplete ring staining with focal condensations (Fig 3B). Furthermore, a sagittal TEM overview revealed CA-induction induced clear morphological changes to the multi-layered polarised cell culture (Fig. 3D). CA-induced spatial alterations to the adherens junction (AJ) protein β-catenin, showing a loss of basal β-catenin (in the apical lining cells of the cell layer), and reduction in apical TJ marker ZO-1 (Fig 3E).

**Figure 3:**
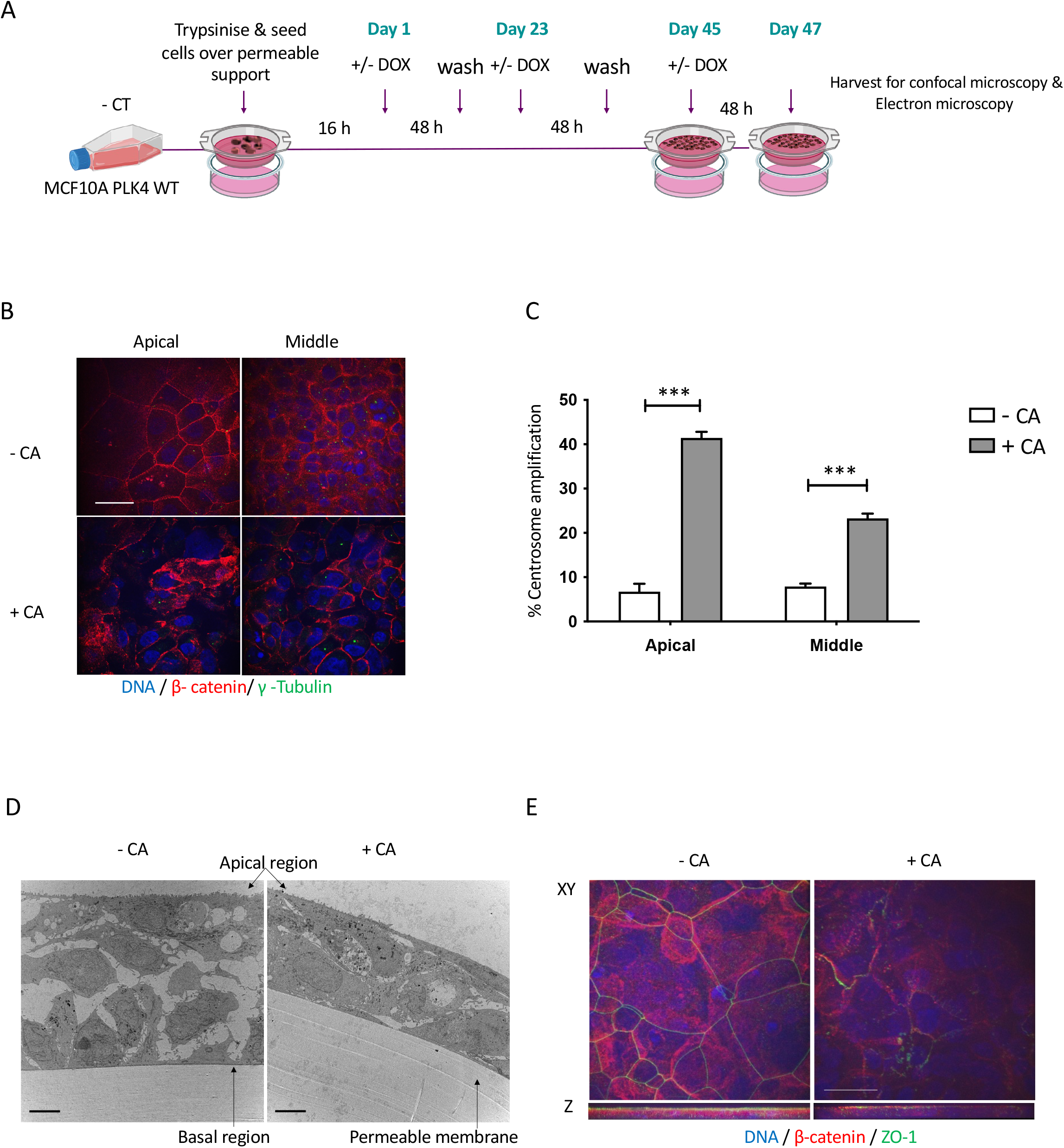
Long-term MCF10A PLK4 WT cell culture transwell system. (A) Experimental layout of long-term transwell membrane cell culture system. (B) Representative immunofluorescent images of MCF10A PLK4WT cells cultured for 47 days. CA was induced by treating at the specified intervals with 2 μg/ml DOX. The cells were fixed and permeabilised in 3.7% PFA & methanol/EGTA. Centrosomes were labelled with g-Tubulin (green) and AJs with β-catenin (red) and DNA counterstained with DAPI. Disrupted and mislocalised β-catenin seen in CA+ cells compared to CA-cells showing well-formed β-catenin at the apical region. Scale bar 20μm. (C) The bar graph shows a significant increase in % CA (***P ≤ 0.001) within the apical and middle layers of the multi-layered long-term culture. Bar graphs represent mean ± SEM from three biological repeats. (D) TEM image of sagittal view of the multi-layered cell structure with altered intercellular space in response to CA induction. Images representative of 4 independent experiments. Scale bar 4μm. (E) XY and Z-stack projections showing disruption of TJ protein ZO-1 (green), AJ protein β-catenin (red) at the apical side in CA+ conditions. Localisation of both ZO-1 & β-catenin was found to be primarily within the apical region rather than the middle or basal regions in long-term culture system. Scale bar 20μm.

### CA disrupts apical junctional complexes

Correct localisation/orientation of cell junction proteins are required for maintaining epithelial cell polarity and epithelial barrier integrity, including TJ proteins JAM-A, occludin and ZO-1, and AJ protein β-catenin. In addition to ZO-1 and β-catenin, CA disrupted TJ family proteins occludin and JAM-A (Fig. 4A, S4A-B). Quantification of cell junction disruption revealed CA induced a significant increase in overall epithelial cell junction disruption (Fig. 4B, p ≤ 0.01, p ≤ 0.001). Quantifying the effects of CA on epithelial barrier intensity/protein distribution there was a significant increase in (*** p < 0.001) β-catenin and JAM-A intensity, and a significant decrease (p ≤ 0.001) in ZO-1 and occludin intensity (Fig. 4C). Detailed analysis of the increased grey value intensity values induced by CA for β-catenin and JAM-A indicated a change to pockets of brighter, more punctate localisation of the protein, whereas ZO-1 and Occludin showed an overall reduction in intensity of staining induced by CA. Graphs represent mean ± SEM from three independent experiments. Confirming these results obtained were specifically due to CA, and not off-target effects of PLK4 overexpression or DOX treatment, PLK4^**1-608**^ overexpression did not cause significant cell junction disruption or protein mislocalisation (Fig. S4C).

**Figure 4:**
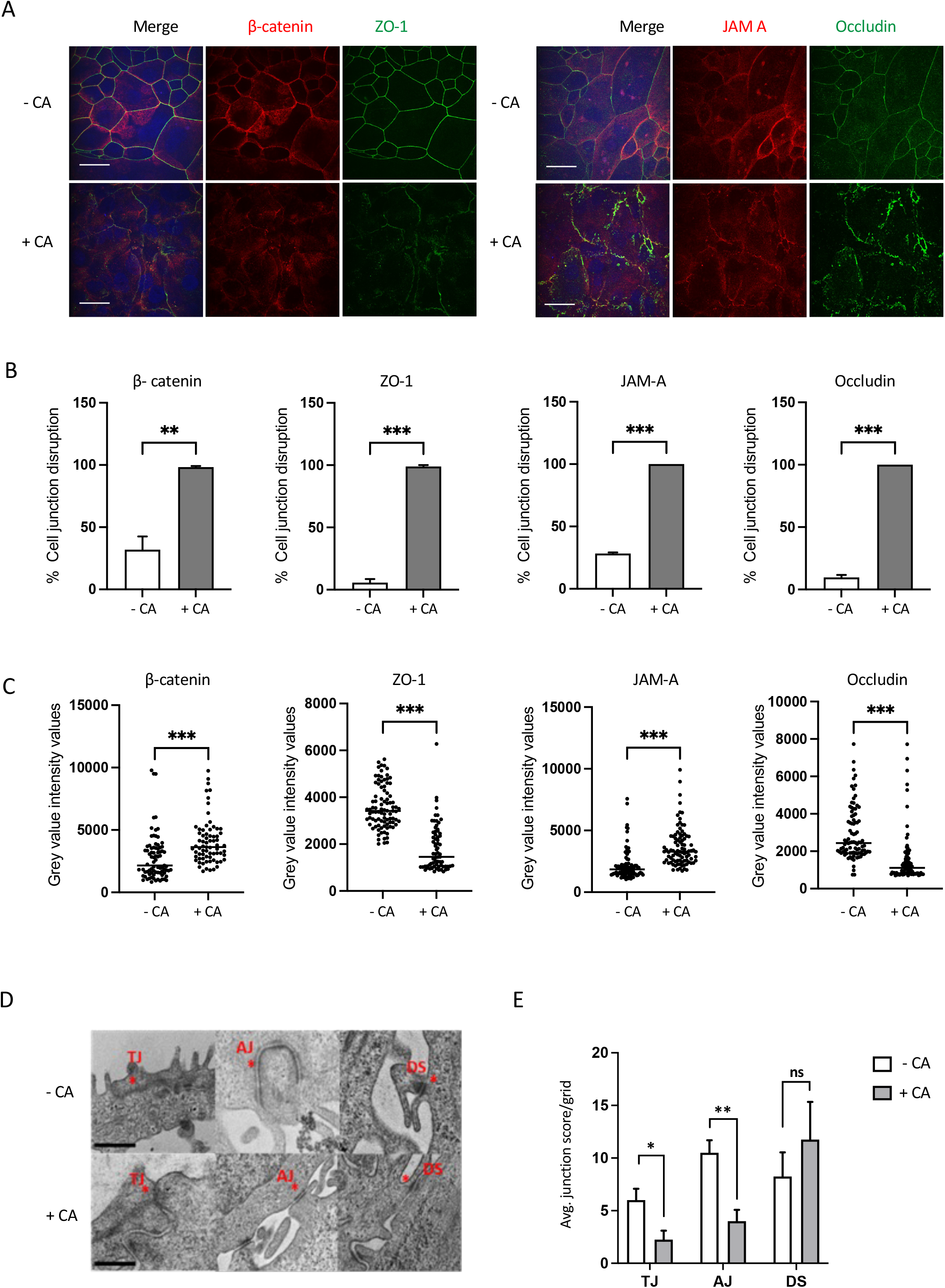
CA disrupts apical junctional complexes in MCF10A PLK4 cells. (A) Representative confocal images of cells co-stained for (left panels) AJ protein β-catenin (red) and TJ protein ZO-1 (green), and (right panels) TJ proteins JAM-A (red), occludin (green) showing localisation and intactness of epithelial cell junctions in CA± MCF10A PLK4 cells. C ± cells were fixed in 3.7% PFA and permeabilised in 0.5% TritonX 100. Images are representative of three biological repeats. Scale bar 20μm. (B) Graph represents % mean ± SEM cell junction staining disruption in 10 fields of view/slide and (C) average grey value intensity values over 10 fields of view. CA significantly (** P ≤ 0.01, *** P < 0.001) disrupted cell junctions and altered the protein localisation. Paired Student T-test was used for was used to determine significance, ** P ≤ 0.01, *** P < 0.001. (D) Representative TEM images of CA± tight junctions (TJ), adherens junctions (AJ) and desmosomes (DS) indicated by asterisk (*) taken at 20000x, HV=100.0kV, Scale bar 600nm. (E) Bar graph represents the cell junction quantification showing a significant downregulation of tight junction and adherens junctions (in apical region) and no significant difference in desmosomes due to CA. Bars represent mean ± SEM junction number from four independent experiments (10 images per experiment), and an unpaired two-tailed student T-test was used to determine the significance, * P < 0.05, ** P ≤ 0.01.

Using TEM analysis, TJs are observed as electron dense apical membrane contact points or “kissing points” where adjacent plasma membranes fuse and form a seal on the intercellular space (Marilyn G. Farquhar & Palade, 1963). The less-electron dense AJs are located immediately below the TJs, also connecting to the actin cytoskeleton, while underlying desmosomes (DSs) connecting to microfilaments appear as electron rich surrounded by a fuzzy area. Ultra-structural investigation of CA effects on cell junction formation, revealed that DSs were the major intercellular junctions bridging inner cells. TJ/AJs were mainly found towards apical region whereas DSs appeared as small intracellular plaques abundant towards the interior and basal layers (Fig. 4D).

Quantifying the effects of CA on cell junctions, CA+ MCF10A had significantly reduced TJs and AJs (p ≤ 0.05, p ≤ 0.01) at the apical side cell-cell interface (Fig. 4E). We noted a non-significant increasing trend in DS numbers in inner cells of CA+ MCF10A grids, suggesting a possible role for desmosomes in cell-cell contact in response to CA disrupted TJ/AJ formation. Furthermore, CA induction induced long bundles of cytoskeletal microfilaments, which were closely associated with intermediate cell junctions (DS), to become acutely truncated and structurally different (undulating) (Fig. 5A). Additionally, CA+ MCF10A consistently displayed increased autophagosome or phagophore-like structures near deformed cell junctions (Fig. 5B-C). Intriguingly, these findings support work by Denu et al, demonstrating that CA disrupts autophagy trafficking by modulating the microtubule network and increases the accumulation of autophagosomes in MCF10A cells (Denu *et al*., 2020).

**Figure 5:**
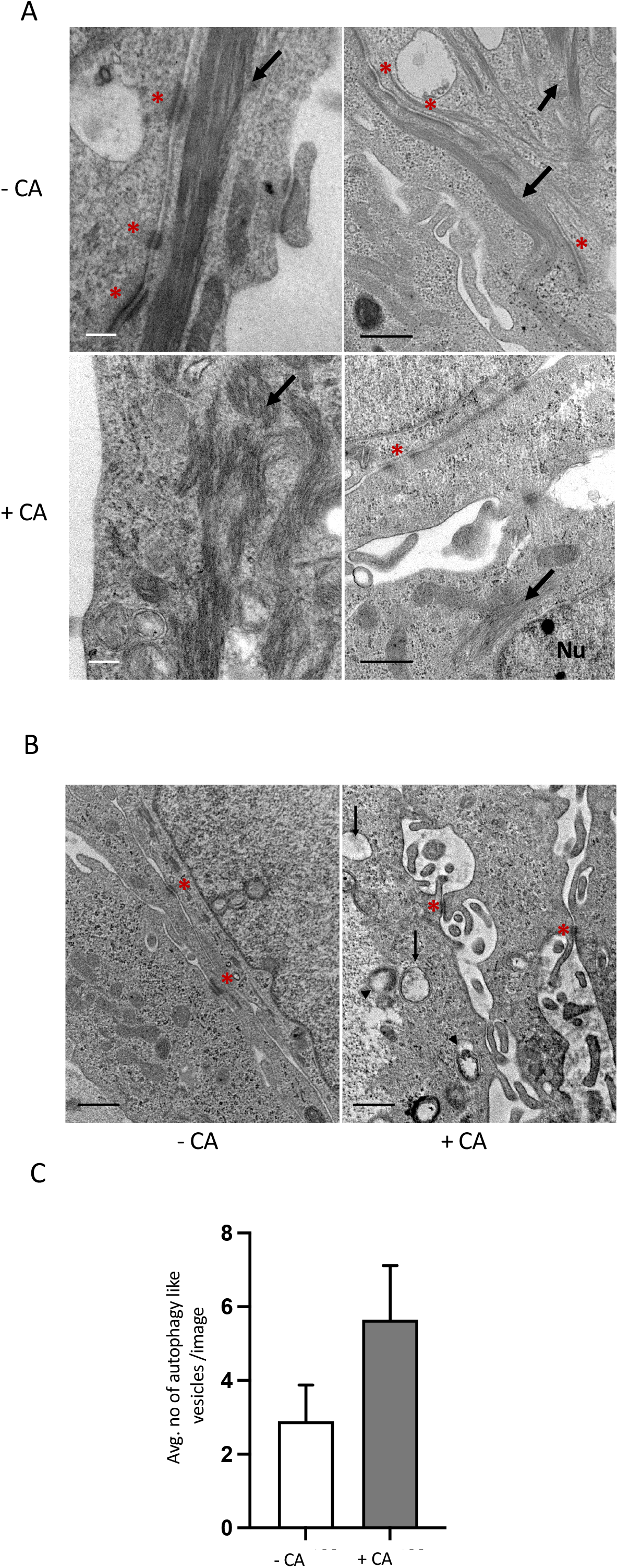
CA alters cytoskeletal structure and increases phagophore and autophagosome-like structures. (A) Representative images of altered microfilament structures in long-term CA+ MCF10A cultures. CA-cells show normal elongated filaments (indicated by arrows) and CA+ show severely truncated, undulating filaments (indicated by arrows). Cell junctions (AJ & DS) are indicated by an asterisk (*), Nu = nucleus. Images were captured at 20000x, HV=100.0kV; Scale bars 200nm (white) and 600nm (black). (B) Representative images of phagophore (arrows) and autophagosome-like (double membrane structure, arrow heads) structures in CA+ cells found proximal to deformed cell junction complexes. DS are indicated by asterisk (*). HV=100.0kV. Scale bar 500nm. (C) Bar graph represent mean (± SEM) number of phagophore/autophagosomes structures observed in CA± in 15 fields of view.

### CA disrupts epithelial cell-cell adhesion in tumourigenic breast cancer cells

To investigate the effects of CA on tumourigenic cells, we chose the luminal A breast cancer cell line MCF7 which assembles (partial) tight junctions under standard long-term culture conditions (Zangari *et al*, 2014). We used the same CA induction system of PLK4 overexpression in MCF7 PLK4 (Arnandis *et al*., 2018). CA was induced in a long-term culture system (Fig. 6A) in MCF7 PLK4 cells (CA+ MCF7, Fig. 6B). In response to CA, quantification of epithelial TJ markers ZO-1, JAM-A and occludin revealed a clear, but non-significant disruption and mislocalisation (Fig. 6C-D). Also, diffusion of β-catenin away from the cortical membrane was observed in CA+ MCF7 (Fig 6C). Ultrastructural analysis on CA+ MCF7 revealed that while CA had no effect on AJ or DS formation, there was a declining trend in TJ formation (Fig 6E-F). Finally, as in MCF10A, CA increased Rap1 activation in MCF7 PLK4 cells (Fig. 6G).

**Figure 6:**
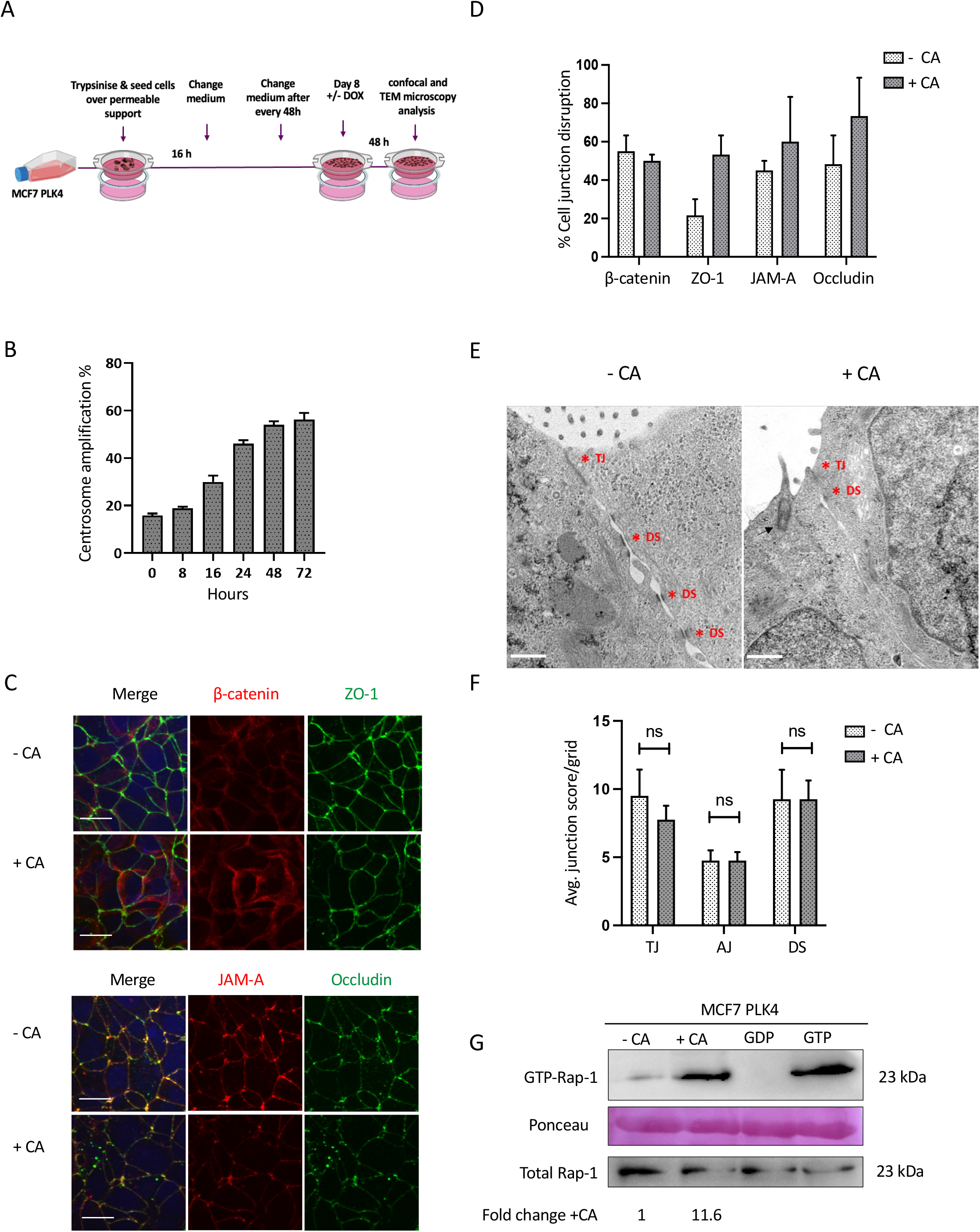
CA disrupts epithelial cell-cell adhesion and increases active Rap1 levels in tumourigenic breast cancer cells. (A) Experimental layout of long-term MCF7 PLK4 cell culture system. (B) Bar graph shows % of CA+ MCF7 PLK4 cells at each time point post-Dox 2μg/ml. Cells were fixed with Methanol/5mM EGTA solution and stained for centrosomes (γ-Tubulin, green), microtubules (α-tubulin, red) and counterstained with DAPI (DNA, blue). Bars represent mean ± SEM, n=3; ≥200 cells/time point. (C) Representative images of CA± MCF7 cells co-stained for AJ protein β-catenin (red), TJ protein ZO-1 (green) (top panels), TJ proteins JAM-A (red) and Occludin (green) (bottom panels). Scale bar 20μm. (D) Bar graph represents % mean ± SEM cell junction staining disruption in 10 fields of view/slide. (E) Representative TEM images of MCF7 PLK4 cells showing cell junctions in ± CA (indicated by asterisk). Cells were imaged at Mag=20,000x, HV=100.0kV, Scale bar 600nm. (F) Bar graph represents cell junction quantification ± CA, with a clear decrease in TJ formation in CA+ MCF7 PLK4 cells. Bars represent mean cell junction number ± SEM from four independent experiments and a two-way ANOVA Bonferroni post-test was performed. No significance was found. (G) Western blot of Rap1 pull-down assay in MCF7 PLK4 shows CA-induced increased GTP-bound Rap-1. Negative (GDP) and positive (GTPγS) controls on untreated cell lysates were included. Ponseau stain is included as a loading control for the pull-down. Total Rap-1 in 60μg of total cell lysates is shown in the bottom panel. The image shown is representative of n=3 independent repeats and the fold-change +CA represents the quantification of ±CA bands by densitometry.

### Profiling early stage CA dysregulation of cell-cell junction and cell-ECM attachment proteins and genes

To map the early stages of CA induction on mechanisms of cell attachment, cell junction and cell adhesion, cell junction total protein expression was evaluated at 48 and 96 hours post-CA induction. A 5-fold reduction in total TJ ZO-1, and 1.4-fold reduction in total TJ occludin was observed in +CA cells (Fig. 7A, upper panels). A ~2-fold increase in total AJ β-catenin and a 1.5-fold upregulation of gap junction connexin 43 (Fig. 7A, middle panels was seen +CA. DS desmoglein 1 displayed a 2-fold increase whereas DS desmocollin was unaffected +CA. Plakoglobin (also known as γ-catenin), an adaptor protein found in both DS and AJ was unaffected (Fig 7A, lower panels). Analysis of subcellular fractions showed that CA induction was associated with redistribution of occludin and β-catenin from the cytoskeletal to the nuclear fractions, with reduced levels of JAM-A in cytoskeletal fraction (Fig. 7B). While ZO-1 levels increased in the cytoskeletal fraction, confocal and TEM analysis demonstrate that this is an abnormal, punctate localisation which does not form a robust cell junction.

**Figure 7:**
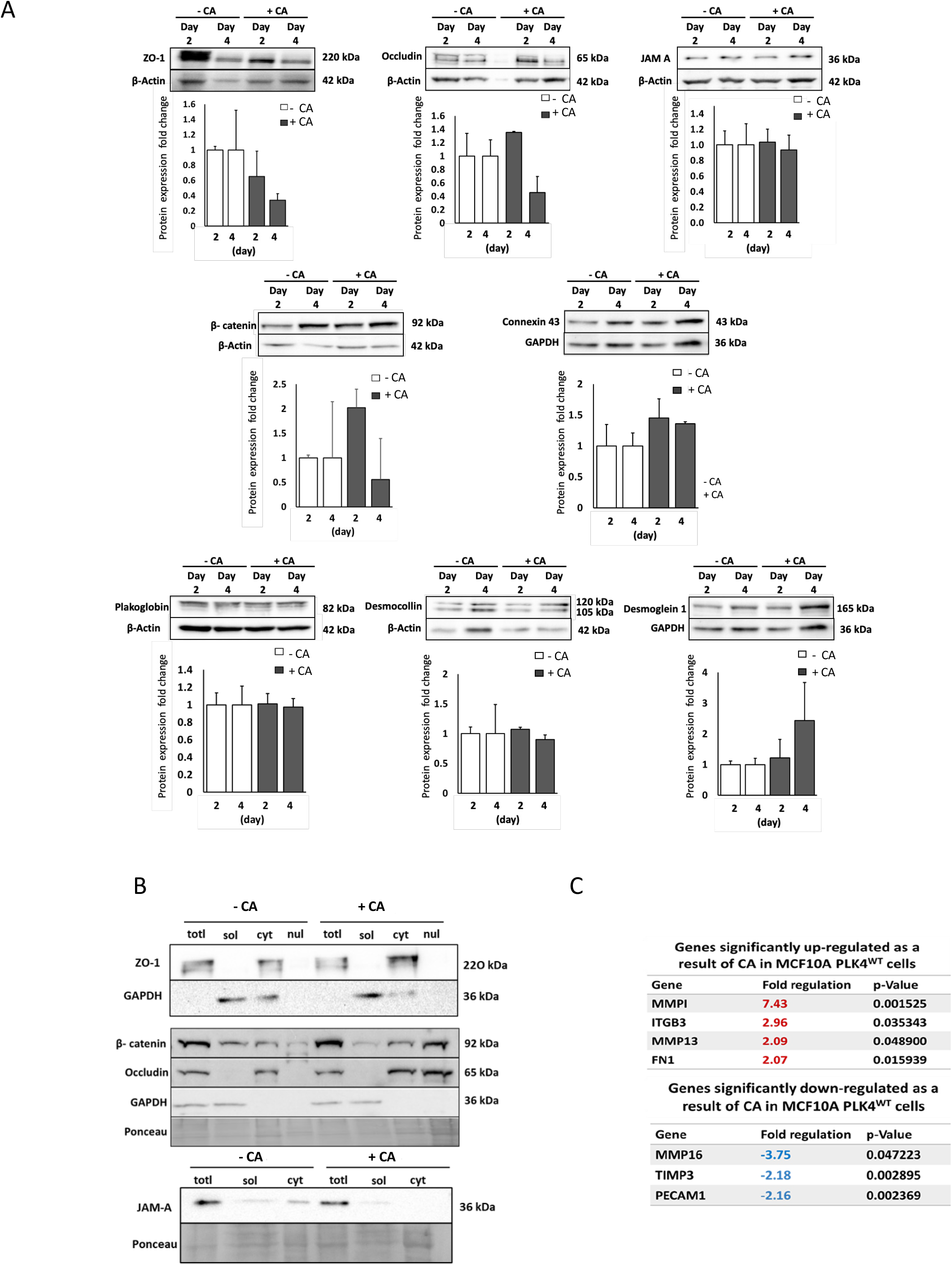
CA dysregulation of cell-cell junction and cell-ECM attachment proteins and genes. (A) Western blot on total protein extracts showing fold-change in expression above β-actin/GAPDH controls (mean±SEM, n=3). (B) Western blot of subcellular fractionation (n=3). Totl=Total, sol=Cytosolic, cyt=Cytoskeletal, nul=Nuclear. CA caused altered recovery of TJ, ZO-1, occludin and JAM-A in cytoskeletal fractions and redistribution of β-catenin and occludin to the nuclear fractions. (C) In RT-PCR analysis of CA± MCF10A, CA significantly upregulated MMP1, MMP13, ITGB3, FN1 and downregulated MMP16, TIMP3 and PECAM1 gene expression. Table shows final data from Qiagen with the CT cut-off value as 35. The p-values were calculated by Student’s t-test of the replicate 2^ (-Delta CT) values for each gene in the control group and treatment groups (n=3 independent experiments).

To further profile dysregulation in response to CA transcriptional analysis of key cell adhesion, cell-ECM gene expression was performed (Fig. 7C). CA induction significantly up-regulated (p = 0.035) the expression of the cell-matrix adhesion molecule expression integrin beta 3 (ITGB3), its binding partner and ECM component fibronectin (FN1) (p = 0.016). ECM degrading enzymes Matrix metalloproteinases were likewise significantly up-regulated, MMP1 (p = 0.001), MMP13 (p = 0.049). MMP16 was significantly down-regulated (p = 0.047), in addition to MMP inhibitor (TIMP3) (p = 0.003) and PECAM1 (p = 0.002).

## DISCUSSION

CA is a known hallmark of cancer detected in a range of human solid and haematological malignancies, particularly associated with increased tumour aggressiveness, poor survival and metastatic cancer subtypes (Chan, 2011; Denu *et al*, 2016). This work supports mounting evidence that CA induction alone can induce tumourigenic transformation of normal cells (Rhys & Godinho, 2017). Here we showed that CA promotes cell migration and invasiveness *in vitro*, independent of the mechanism of induction (PLK4 overexpression or CDK1 inhibition). This work quantifies the dissemination of CA+ pro-tumourigenic cells out of the epithelial layer via a combination of cell autonomous and non-cell-autonomous effects of CA previously described (Arnandis *et al*., 2018; Godinho *et al*., 2014). This was supported by *in vivo* validation (using the Chick embryo xenograft model (Harper *et al*., 2021)) which demonstrated the earliest possible observable effects of CA induction, with cell invasion into the chicken mesoderm, hyperplasia of the chorionic epithelium at the interface with the Matrigel-cell graft, heterophil infiltrates (the avian heterologues of neutrophils) and dilated lymphatics. Additional secondary tumour-associated characteristics of CA induction found *in vivo* included hemorrhaging, central necrosis and calcification. This clearly demonstrates that induction of CA alone, in a control non-tumourigenic background, is sufficient to trigger extensive pro-tumourigenic characteristics within days. These findings further inform the role of CA in tumourigenesis observed using mouse models (Coelho *et al*., 2015; Levine *et al*., 2017; Sercin *et al*., 2016).

Signalling pathways involving small GTPases, Rho and Ras, frequently shown to be dysregulated in tumours, have previously been proposed to mediate CA-induced cellular effects. The Rho subfamily consisting of Rho A, Rac1, Cdc42 is well known for regulating cytoskeleton and actin dynamics, and Rac1 has been associated with lamellipodia formation during cell migration (Sit & Manser, 2011). Increased microtubule nucleation by supernumerary centrosomes upon CA induction leads to increased Rac1 activity which promotes actin polymerisation disrupting cell adhesion and promoting cell invasion (Godinho *et al*., 2014). In addition, small GTPase Ras-associated protein Rap1 is highly expressed in cancer cells, stimulating integrin mediated cell-ECM attachment, matrix metalloproteinase expression involved in ECM remodelling and cytoskeletal modulation, paving the way for cancer cell invasion and migration (Zhang *et al*, 2017). We found that CA induction increases active GTP-bound Rap1, and that CA-dependent cell-ECM attachment and invasion is highly dependent on this Rap1-mediated signalling. *In vivo*, Rap-1 inhibition blocked key tumourigenic phenotypes displayed by centrosome amplified cells (invasion, hyperplasia, and dilation of lymphatics and blood vessels). Therefore, in addition to Rac1, Rap1 GTPase plays a key role in the signalling pathways mediating CA induction of an invasive phenotype in control breast epithelial cells and this points to Rap1 inhibition as a potential new therapeutic option for many CA tumours. Rap1 acts as a tumour promoter mediating metastasis in a range of cancer contexts including breast, prostate, ovarian, oesophageal and pancreatic cancers (reviewed in (Looi *et al*, 2020)) and this work informs the mechanism by which Rap1 works.

Our work shows that CA induction in an untransformed epithelial background is sufficient to trigger migration and invasion *in vitro* and *in vivo*, in conjunction with elevated cell-ECM attachment. Previous work has shown that tumourigenesis is associated with dramatic alterations in integrin expression profiles with these adhesion molecules facilitating cell movement by forming focal adhesions between cancer cells and the ECM (reviewed in (Hamidi & Ivaska, 2018). In breast cancer, integrin β3 overexpression (bound to subunit αv to form αvβ3 integrin) is associated with high metastatic potential, especially in promoting bone metastasis (Carter *et al*, 2015; Havaki *et al*, 2007; Kovacheva *et al*, 2021). Integrin β3 overexpression in breast epithelia paves the way for invading cells by inducing secretion of matrix metalloproteases MMP1 and 2 with resultant matrix degradation. Fibronectin binding to integrin β heterodimeric receptors stimulates the production of MMP-1 inducing an MMP-1-dependent invasive phenotype in breast cancer cells (Jia *et al*, 2004). Our data show CA-induced upregulation of gene expression of ITGB3 alongside that of its ECM binding partner fibronectin, and dysregulation of the balance of ECM degrading enzymes matrix metalloproteinases (MMP1, MMP13 and MMP16) and Tissue inhibitors of MMPs (TIMP1). These effects of CA support clinical studies reporting high fibronectin expression in metastatic breast cancer is associated with high MMP expression (Das *et al*, 2008; Fernandez-Garcia *et al*, 2014).

The formation of apico-basal polarised epithelia is closely coupled to cell-cell adhesion events at apical junctional complexes (AJC) consisting of TJ and sub-adjacent AJ, helping regulate normal cell polarity and movement and inhibiting abnormal migration and metastasis (Wang & Margolis, 2007). TJs act as a semi-permeable adhesive barrier regulating traffic through the paracellular space (gate function) while physically segregating lipids between apical and basolateral compartments. AJs contribute to cell-cell adhesion via an adhesive belt lying basally to TJs. The cytoplasmic domains of the proteins that comprise AJC junctions are tethered to the cytoskeleton via various adaptor proteins integrating signals to regulate cell polarity and movement. We observed that CA disrupted and mislocalised TJ and AJ components in MCF10A and MCF7 cell lines. The subcellular mislocalisation of JAM-A, occludin and β-catenin away from the cytoskeleton as well as overloading of ZO-1 protein at the cytoskeleton in CA+ cell populations, is a likely underlying cause of disrupted of cell-cell adhesion. It is likely that excessive centrosome numbers impact on the cytoskeleton protein dynamics via elevated microtubule nucleation followed by actin polymerisation (Godinho *et al*., 2014). Intriguingly, microtubule polymerisation inhibitors (MTIs) can re-establish functional AJCs in cell-cell adhesion-defective carcinoma cells via mechanosensitive recruitment of junction molecules (Ito *et al*, 2017). In this case, GTPase RhoA was found to be upregulated by MTIs, restoring AJCs by causing actomyosin contraction at the apical cortex and tension-dependent junction reassembly. The possibility that the opposite effect may be happening in our CA+ cells with elevated Rap1-GTP regulating RhoA activity remains to be explored.

The treatment of transformed cancer cell lines with PLK4 inhibitor Centrinone B to decrease inherent CA and reverse CA-induced phenotypes has previously been demonstrated (Denu *et al*., 2020; Denu *et al*., 2018; Wong *et al*., 2015). In the current study, differential effects Centrinone B on invasive capabilities of the cells correlated with endogenous CA levels, where the confirmed contribution of CA to migration and invasion of triple negative MDA-MB-231 breast epithelial cells (high CA) was not recapitulated in MDA-MB-468 (low CA). This indicates that across cancer cell subtypes, there are varying contributions from CA and other cellular mechanisms driving invasive capacity.

In conclusion, we found that CA induces tumorigenic changes in breast epithelial contexts *in vitro* and *in vivo*, promoting the transition to an invasive phenotype via disruption of cell-cell contacts and release of stromal proteases to facilitate destruction of the basement membrane and increased motility of invasive cells through the ECM. The *in vivo* finding that CA is sufficient to confer such advanced tumourigenic changes within days, identifies CA as an early driver alteration in cancer and a central targetable mechanism.

## Abbreviations

CA: Centrosome amplification
CAM: Chicken Chorioallantoic membrane
MTOC: microtubule organising centre
NLP: ninein-like protein
HUVEC: human umbilical vein endothelial cells
CA+: CA-induced
DOX: Doxycycline
IF: Immunofluorescence

## ACKNOWLEDGMENTS

We thank Susana Godinho, Barts Institute, London for the MCF10A PLK4 and MCF7 PLK4 cell lines and Maura Grealy, University of Galway for antibodies. The authors would like to acknowledge the Hardiman scholarship, University of Galway, CMNHS Millennium fund and the Irish Association for Cancer Research Education Grant Award for financial support. SHV was supported by Science Foundation Ireland grant 13/IA/1994 (to AMH). Imaging in this project was carried out in the Centre for Microscopy & Imaging at the University of Galway (www.imaging.nuigalway.ie), a facility that is funded by the Irish Government’s Programme for Research in Third Level Institutions, Cycles 4 and 5, National Development Plan 2007-2013.

## DECLARATION OF INTERESTS

The authors report no conflicts of interest.

## MATERIALS AND METHODS

### Cell culture

Wild type MCF10A (ATCC) and MCF10A PLK4, MCF10A^1-608^ doxycycline-inducible PLK4 cell lines (a gift from Susana Godinho, Barts Institute (Godinho *et al*., 2014)) were used to model CA in an untransformed breast cell context following 48h doxycycline at 2 mg/ml. MCF10A were cultured in Dulbecco’s Modified Eagles’ Medium/F-12 media supplemented with 5% Horse serum (Sigma 14M175/ H1270), 0.5μg/ml Hydrocortisone (Sigma H0888), 20ng/ml human Epidermal Growth Factor (hEGF) (Sigma E9644), 10μg/ml Insulin (Sigma I9278) 1% Penicillin/Streptomycin (Sigma P4333). The breast cancer cell lines MDA-MB-231, MDA-MB-468 and doxycycline-inducible MCF7 PLK4 cells (a gift from Susana Godinho, Barts Institute (Arnandis *et al*., 2018)) were cultured in RPMI-1640 (Sigma R8758) supplemented with 10% Fetal Bovine Serum (Sigma F7524) and 1% Penicillin/Streptomycin (Sigma P4333). For CA induction, cells were treated with 2μg/ml Doxycycline hyclate (Sigma D9891) or 10μM CDK1 inhibitor, RO3306. 150nM Centrinone B treatment for 16 h inhibited CA. Rap1 was inhibited with 10 μM GGTI-298 (Biotechne Tocris 2430) for 3 h.

### Immunofluorescence microscopy

Cells were seeded on Poly-D-Lysine (Sigma P7280) coated coverslips and fixed in 2 ml chilled Methanol/EGTA (5mM) (Sigma E4378) at −20°C. Cells were incubated with primary antibodies against centrosome (γ-tubulin Sigma T3559) and microtubule (α-tubulin Sigma B512) markers and stained with secondary antibodies (AlexaFlour^®^ 594, Invitrogen A-11005, AlexaFlour^®^ 488, anti-rabbit Invitrogen A-11008). Coverslips were mounted in Vectashield with DAPI (Vector Laboratories H-1200), sealed with nail varnish and stored at 4°C. Imaging was performed on the OLYMPUS BX61 fluorescent microscope driven by Olympus cellSens imaging software and Z-stacks (0.4μm) captured. Centrosome number was quantified in ≥200 cells per slide categorising normal as 1-2 centrosomes/cell or centrosome amplification as ≥3 centrosomes.

### Cell invasion and migration assays

CA was induced for 36h with 2 μg/ml DOX in MCF10A PLK4 then cells serum-deprived (1% horse serum and no hEGF) overnight (12h). Cells were seeded on transwell inserts (8-μm pore, Corning 353097) - coated with matrigel (Corning™ 356231) for invasion assay; uncoated for migration assay. To create a gradient, complete media + 6 % horse serum and 20 ng/ml hEGF was added to the lower chamber. For negative gradient controls, serum-deprived media (1% horse serum and no hEGF) was added to the upper and lower chambers. Cells were incubated for 12 h and 24h to measure cell migration and invasion respectively. Cells on the underside of the filter post-incubation were stained and quantified using crystal violet (Sigma HT901-8FOZ). Brightfield images from five fields of view per insert were captured on the OLYMPUS CKX31, under 20X magnification. Migrated and invaded cells were quantified using ImageJ software wand tool (Schneider *et al*, 2012).

### Chicken Embryo Xenograft Model

Fertilised chicken eggs (Breed: Ross 308, 3 eggs per treatment) were incubated at 37°C/ 65% humidity. On embryonic development (ED) day 3, a window was opened in the eggshell under a heat lamp and 2.5ml albumen aspirated off using a syringe. The eggs were re-sealed with transparent adhesive film (Opsite Flexifix™, Smith-Nephew). On ED day 8, MCF10A PLK4 ± CA ± 10 μM GGTI-298 in 1X PBS:Matrigel (Corning: 354248; 50:50 ratio) was implanted onto the chicken membrane within a sterile silica ring. On ED day 13, the embryos were sacrificed, and the cell-matrigel graft with attached chorioallantoic membrane (CAM) harvested. Gross images were captured using a stereo microscope (Leica S6E), and the tissue fixed and paraffin-embedded in the Thermo Scientific™ Excelsior Tissue Processor unit. 5μm tissue sections were prepared using a Microtome (Microm HM325, Thermo Scientific), mounted and stained with haematoxylin and eosin stain (H&E) and imaged as Z-stacks by brightfield microscopy at 40x on the Olympus VS120 Virtual Slide Scanner Microscope.

### Immunohistostaining of xenograft tissue

The Cytokeratin 8 monoclonal antibody, (Cytokeratin CAM 5.2 BDBiosciences-345779) was used to label MCF10A cell invasion into the chicken membrane. Slides were deparaffinised, blocked and incubated with anti-Cyt8 antibody, HRP-conjugated secondary antibody followed by 3,3’-Diaminobenzidine (DAB) staining using the ImmPRESS™ Excel Amplified HRP peroxidase Polymer Staining Kit (Vector Laboratories MP-7602-15), according to the manufacturer’s protocol. All images were captured as Z-stacks of brightfield microscopy at 40X magnification on the Olympus VS120 Virtual Slide Scanner Microscope.

### Long-term transwell cell culture

Long-term cell culture in MCF10A PLK4/^**1-608**^ cells was adapted from (Marshall *et al*., 2009). MCF10A PLK4/^**1-608**^ cells cultured in complete DMEM/F12 medium in the absence of cholera toxin (CT) were seeded onto the transwell inserts (1.2×10^5^ cells, pore size 0.4 μm). The cells were maintained by replacing media in both chambers every 48h. Previous studies reported that CA prevalence is higher in *in vivo* tumours than in *in vitro* cells due to intratumoural hypoxia, and that cells with high CA levels *in vitro* are eliminated over time (Mittal *et al*, 2017). To maintain levels of CA, MCF10A were pulsed with Dox (2μg/ml) at regular intervals (Day 1, 23, 45) of the long-term culture (Fig. 3A).

MCF7 PLK4 cells were analysed in a 10-day transwell cell culture system (Fig. 6A) adapted from (Sheller *et al*, 2017).

Transwell inserts were sliced into quarters with a sharp scalpel blade and mounted onto Superfrost slides. Cells were fixed with 3.7% paraformaldehyde and permeabilised with 0.5% Triton X in PBS and hybridised with cell junction primary antibodies JAM-A (sc-53623), β-Catenin (sc-7963), Occludin (Invitrogen 71-1500), ZO-1 (Invitrogen 61-7300) and secondary antibodies (AlexaFlour^®^ 594 Invitrogen A-11005, AlexaFlour^®^ 488 antirabbit Invitrogen A-11008). Filters were mounted with Vectashield + DAPI and images captured with an Andor revolution spinning disk confocal microscope under 60X magnification. The cell junction integrity on the apical side was quantified using binary scoring method, 1 = intact cell junction and 0 = disrupted cell junction. Cell junction protein localisation was quantified using the value intensity (maximum intensity projection of stacks converted to greyscale) of cell junctions marked by β-Catenin, ZO-1, JAM-A and Occludin, at the apical region for 10 fields of view, using ImageJ FIJI Broadly Applicable Routines-(BAR) plugins.

### Transmission electron microscopy (TEM)

Long-term cultures of MCF10A PLK4/^1-608^ and MCF7 PLK4 seeded on filter inserts were fixed in 2% glutaraldehyde (Sigma G5882), 2% paraformaldehyde (Fisher Scientific 10131580) in 0.1M sodium cacodylate pH 7.2 (Sigma C0250). Filters were washed in 0.2M sodium cacodylate buffer, incubated in 1% osmium tetroxide in 0.1M sodium cacodylate buffer (pH 7.2). Filters were dehydrated through a graded series of ethanol concentrations and infiltrated with resin:acetone mixtures overnight (50:50 mixture, 75:25 mixture and 100% resin). Ultra-thin (70-90nm) sections were cut in a sagittal plane using a diamond knife and mounted onto 3mm copper grids. Grids were counterstained with UA Zero non-radioactive EM Stain (Agar Scientific AGR1000) and imaged on a Hitachi 7500 TEM microscope under 20,000X magnification. Cell junctions were quantified in 10 images per grid (4 grids per treatment in 4 independent experiments) and a binary score of 1 and 0 assigned to presence or absence of each cell junction respectively.

### Immunoblotting

Whole cell lysates were prepared as previously described in (Gao *et al*, 2014). A subcellular protein extraction was performed as described in (Choi *et al*, 2014). Briefly, cells were lysed in 50mM PIPES, 50 mM NaCl, 5% Glycerol, 0.1% NP-40, 0.1% Triton X-100 and 0.1% Tween 20 with a protease inhibitor cocktail. The supernatant (cytoplasmic fraction) was retained, and cells rinsed in 5 ml of 50mM Tris-HCl (pH 7.4) and nuclease buffer containing 10 U/mL Benzonase^®^ Nuclease (Sigma E1014), 10 mM MgCl2 and 2 mM CaCl2 in 50 mM Tris-HCl buffer, pH 7.5. The supernatant was retained (nuclear fraction), and cytoskeletal proteins collected by solubilising the pellet in 0.1% SDS.

The protein concentration was determined by Pierce™ Bicinchoninic acid (BCA) Protein Assay according to the manufacturer’s instructions. Western blotting was carried out with standard protocols, 30 μg and 60 μg protein were used for semi-dry and wet transfer respectively. Antibodies used were JAM-A sc-53623, β-Catenin sc-7963, Occludin Invitrogen 71-1500, ZO-1 Invitrogen 61-7300, Plakoglobin BD 61025, Desmoglein-1 BD 610273, Desmocollin 2/3 Zymed 32-6200, Connexin 43 Proteintech 15386-1-AP, anti-Actin Sigma-Aldrich A2066, anti-GAPDH Sigma-Aldrich G9545, HRP-conjugated anti-mouse and rabbit - Jackson ImmunoResearch Laboratories Inc.

### Rap-1 Activation Pull-down Assay

MCF10A or MCF7 cells were seeded in 150mm dishes and CA was induced with 2μg/ml Dox for 48 hours. The assay was performed using the Rap1 Activation Assay Kit (17-321 Sigma-Aldrich) according to the manufacturer’s instructions.

### RNA preparation and real-time quantitative PCR (qPCR)

The Qiagen RT2 Profiler PCR Array for 84 Cell Adhesion and Extracellular Matrix molecules (PAHS-013Z) was used and the PCR component mix for 96 well plate prepared according to manufacturer’s instructions. The experiment was performed as per the Profiler array format C protocol on a StepOnePlus™ Real-Time PCR System (Applied Biosystems). Data analysis was carried out using the Qiagen GeneGlobe RT-qPCR Data Analysis web tool.

### Cell adhesion assay

Assay was performed as previously described in (Chen, 2012). Briefly, 48 h of CA induction using 2μg/ml DOX and overnight serum starvation, cells were transferred to type I collagen (Corning™ 354231) (positive control) or 1% BSA (negative control) coated 96 well plates and incubated for 20 min. After a series of washing steps, the cells bound to the substrate were quantified by crystal violet staining and the ImageJ software wand tool (Schneider *et al*., 2012).

### Statistical analyses

All statistical comparisons were performed with GraphPad^®^ Prism 9 (www.graphpad.com). Depending on the number of factors, multiple-group comparisons were made using either a single or two-way analysis of variance (ANOVA) followed by the Bonferroni test to confirm statistical differences between groups. Paired sample analysis was performed using a two-sided unpaired or paired Student’s t-test. Statistical significance (p-value) was assigned for values <0.05.

## SUPPLEMENTAL FIGURE LEGENDS

**Figure S1**: **CA induced by CDK1 inhibition increases *in vitro* migration and invasion and blocking CA does not effect metastatic characteristics of MDA-MB-468.** (A) Bar graph representing % cells +CA in WT MCF10A post CDK1 inhibitor RO3306 (10 μM) treatment. Bars represent mean ± SEM from 3 independent experiments, ≥200 cells/time point. (B) CA increases cellular invasion and (C) migration in WT MCF10A. (D) Blocking CA using Centrinone B in TNBC MDA-MB-468 cells (with low endogenous CA) did not significantly effect migration and invasion.

**Figure S2: Representative images of chicken xenograft model ± CA. (A-C)** Further representative images of the excised CA± MCF10A-matrigel grafts onto the chicken chorioallantoic membrane (n=4). +CA grafts show marked reactions of the chick chorioallantoic membrane to the matrigel-cell graft with hyperplasia and inflammatory infiltrate. In (A) and (C), the +CA sample shows vascular proliferation and infiltration of the CE into the matrigel layer. Zero or a mild reaction was observed in the -CA grafts.

**Figure S3: CA induces early secondary tumour characteristics in chicken xenograft model** (A) Gross images of excised CA ± MCF10A PLK4/matrigel graft captured on a stereo microscope (Leica S6E). (B) CA+ cell engraftment causes multifocal moderate to marked reaction of the chick chorioallantoic membrane to the Matrigel in contrast to a zero to mild reaction seen in the CA-graft. (C) The CA+ sections (enlarged box) show increased epithelial hyperplasia, inflammatory infiltrate and secondary tumour characteristics including vascular proliferation, haemorrhaging with accompanying central necrosis.

**Figure S4**: **Doxycycline treatment and overexpression of truncated PLK4^1-608^ variant has no effect on localisation of tight junction and adherens junction proteins.** Additional representative images long-term cell cultures. (A-B) Immunofluorescent co-staining of MCF10A PLK4 ±CA shows combinations of AJ protein β-catenin (red), TJ protein ZO-1 (green), and TJ proteins JAM-A (red), occludin (green). Images are representative of 4 independent biological repeats. Scale bar 20μm. (C) Representative images of long-term cell cultures of negative control MCF10A^**1-608**^ ± Dox co-stained for AJ protein β-catenin (red), TJ protein ZO-1 (green) and TJ proteins JAM-A (red), occludin (green) (n=2). Scale bar 20μm. Doxycycline treatment and overexpression of PLK4^**1-608**^ did not disrupt apical junction complex protein localisation.

## Notes

### Competing Interest Statement

The authors have declared no competing interest.

### Summary of Updates

Figures 1-6 and S2 have been revised based on reviewers comments.

