## Supplementary figures and images for "“Centrosome Amplification promotes cell invasion via cell-cell contact disruption and Rap-1 activation”"

### Supplemental Figs S1-4

FIGURE S1

Prakash et al, 2022

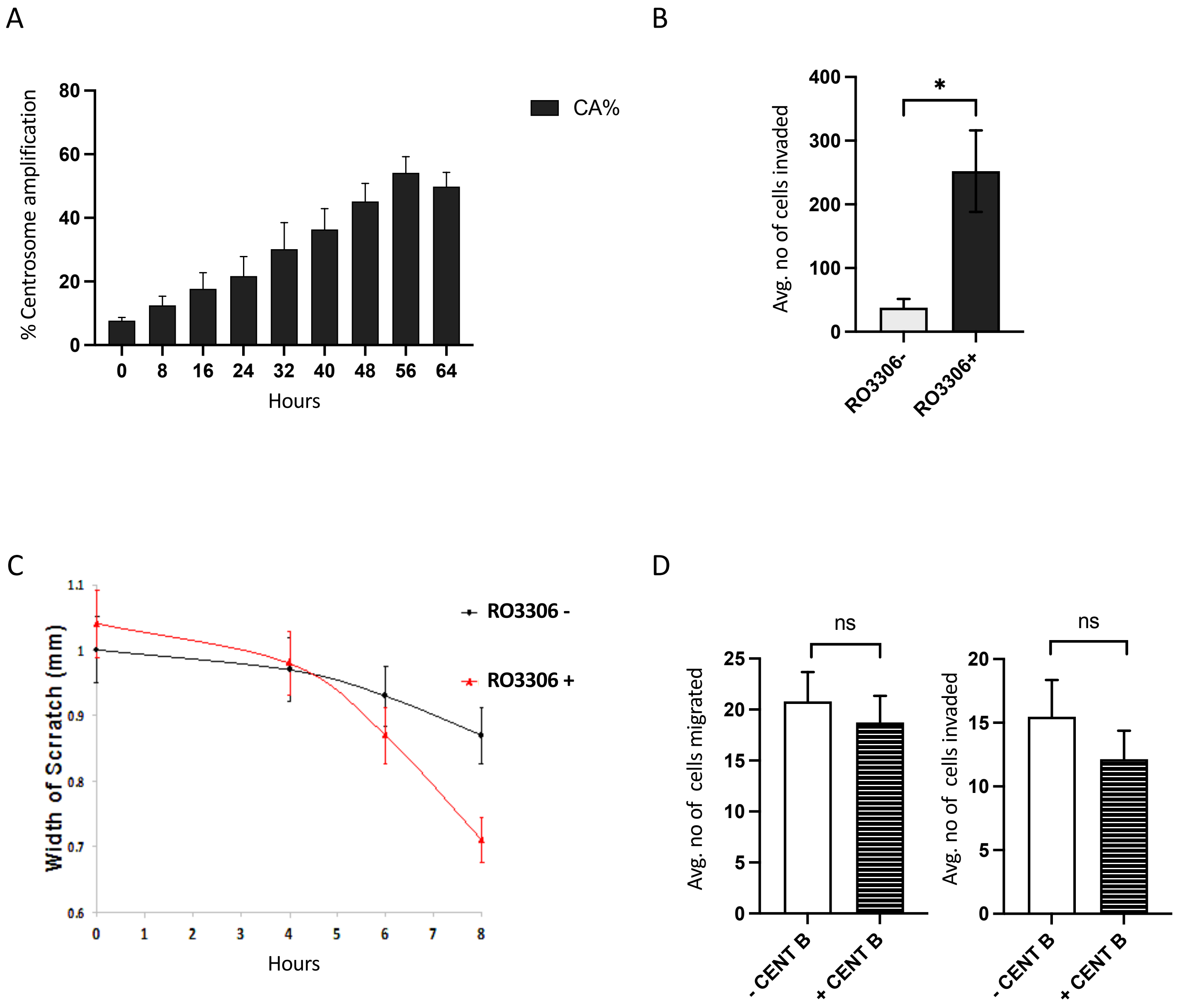

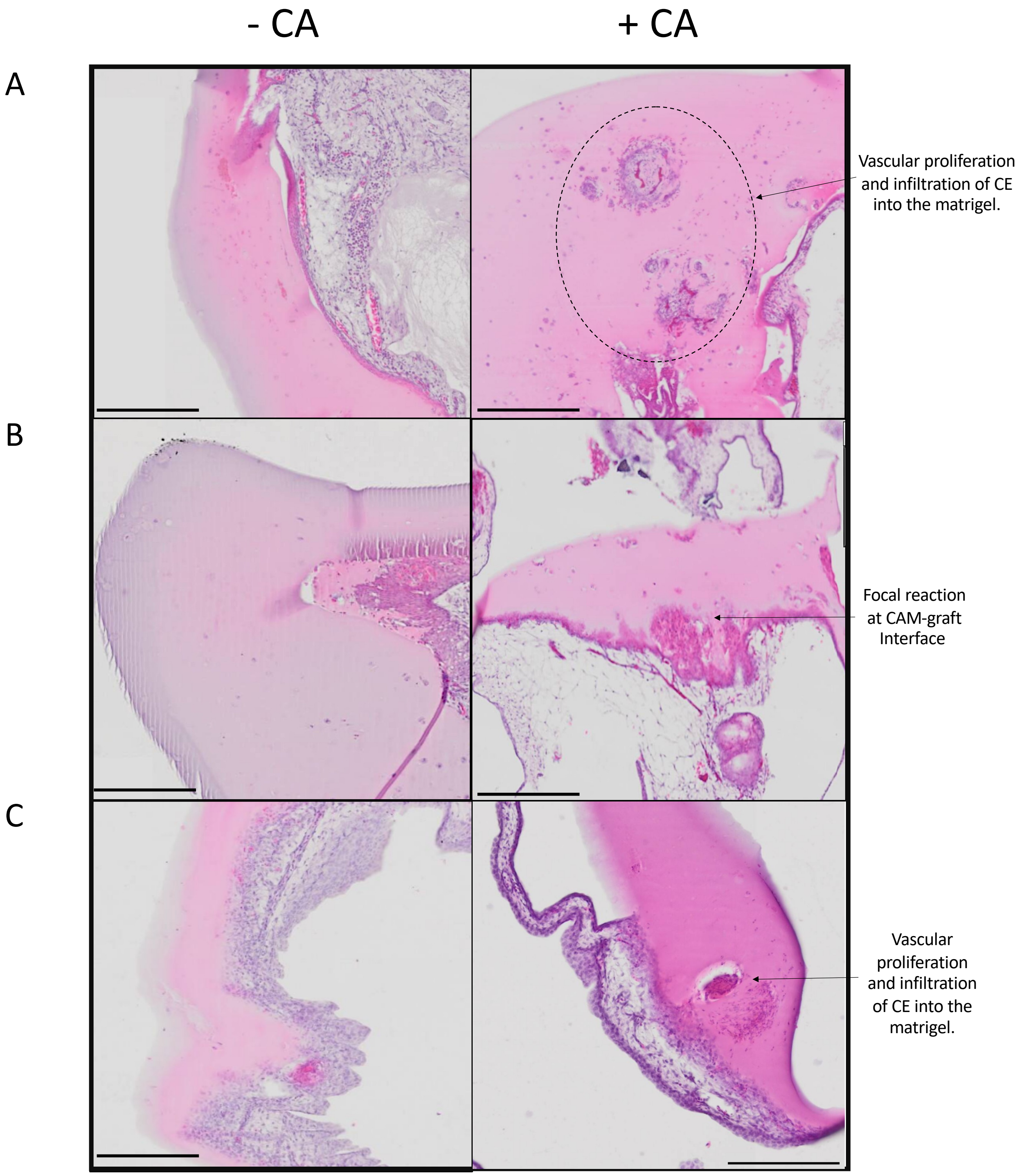

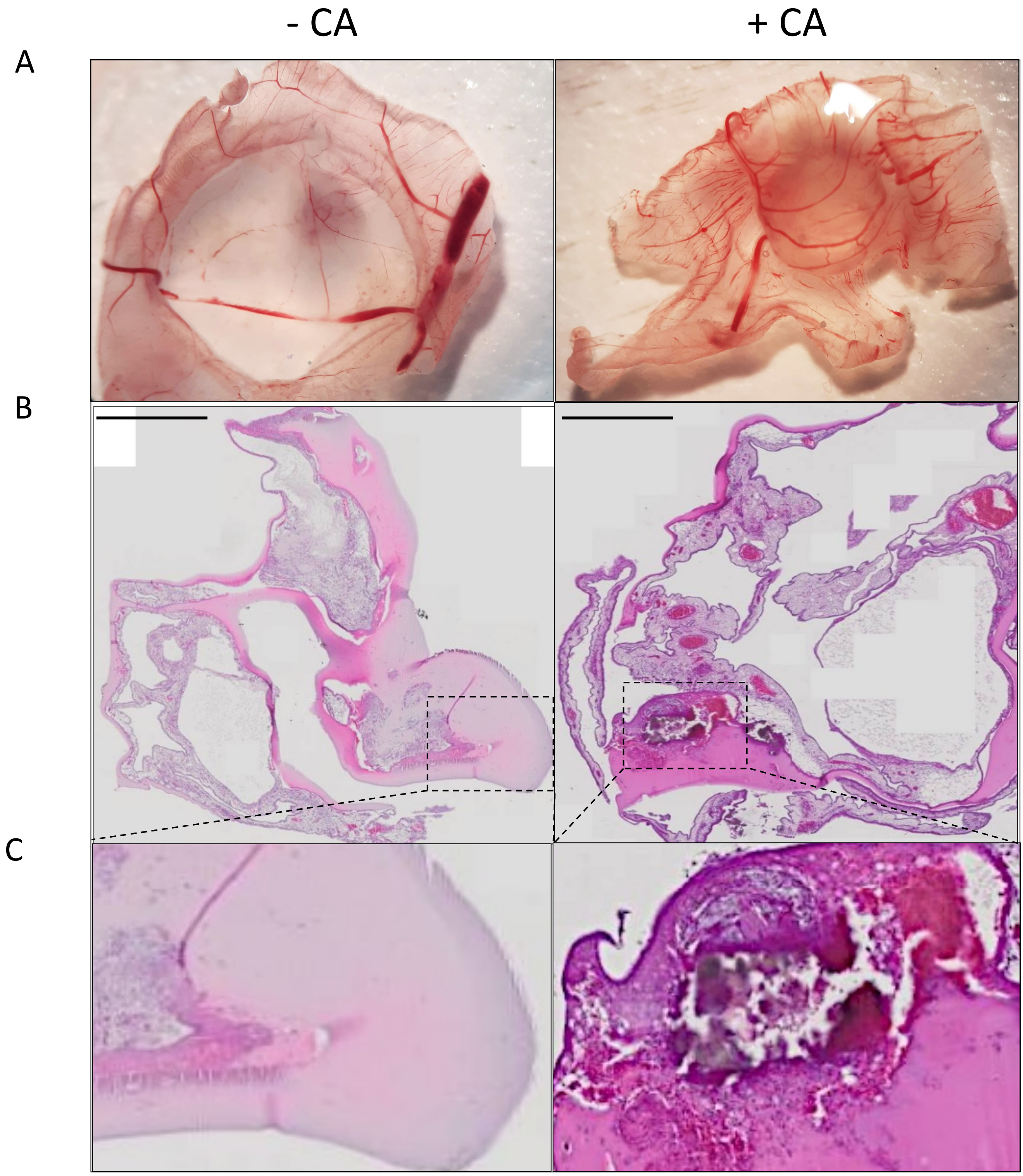

#### FIGURE S4

Prakash et al, 2022

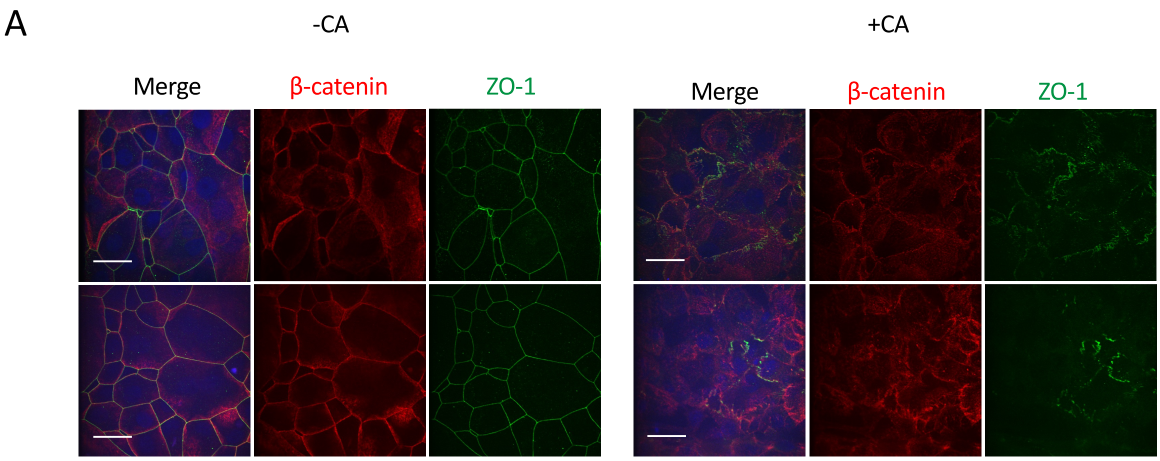


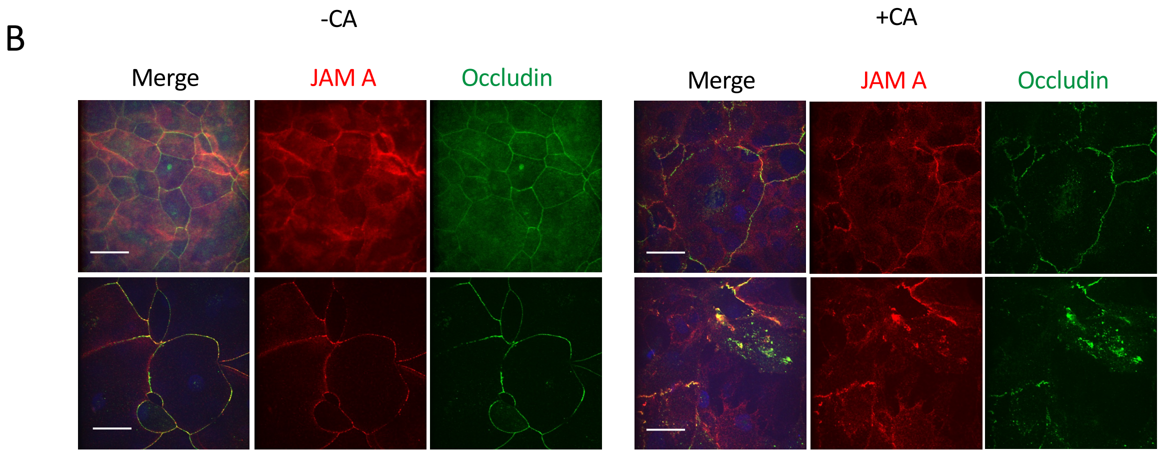


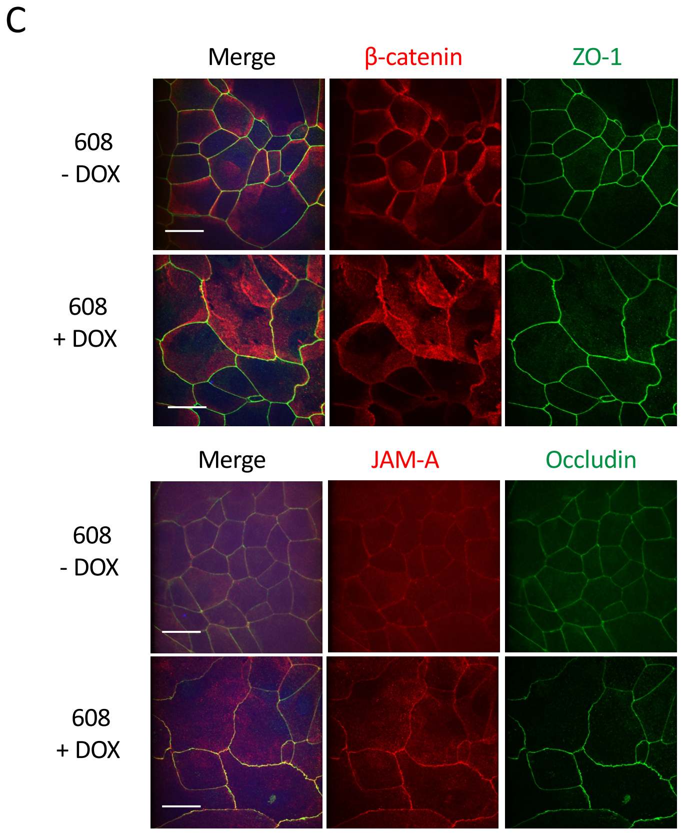
